# Slice-direction geometric distortion evaluation and correction with reversed slice-select gradient acquisitions

**DOI:** 10.1101/2021.12.16.472821

**Authors:** Anna I. Blazejewska, Thomas Witzel, Jesper L.R. Andersson, Lawrence L. Wald, Jonathan R. Polimeni

**Author notes:** Correspondence should be directed to: Anna Blazejewska, Ph.D., MGH Athinoula A. Martinos Center for Biomedical Imaging, Bldg 149 13^th^ St Rm 2301, Charlestown MA, 02129 USA.

## Abstract

Accurate spatial alignment of MRI data acquired across multiple contrasts in the same subject is often crucial for data analysis and interpretation, but can be challenging in the presence of geometric distortions that differ between acquisitions. It is well known that single-shot echo-planar imaging (EPI) acquisitions suffer from distortion in the phase-encoding direction due to B_0_ field inhomogeneities arising from tissue magnetic susceptibility differences and other sources, however there can be distortion in other encoding directions as well in the presence of strong field homogeneities. High-resolution ultrahigh-field MRI typically uses low bandwidth in the slice-encoding direction to acquire thin slices and, when combined with the pronounced B_0_ inhomogeneities, is prone to an additional geometric distortion in the slice direction as well. Here we demonstrate a presence of this slice distortion in high-resolution 7T EPI acquired with a novel pulse sequence allowing for the reversal of the slice-encoding gradient polarity that enables the acquisition of pairs of images with equal magnitudes of distortion in the slice direction but with opposing polarities. We also show that the slice-direction distortion can be corrected using gradient reversal-based method applying the same software used for conventional corrections of phase-encoding direction distortion.

## Introduction

Many neuroimaging studies using MRI frequently leverage multiple imaging protocols with varying complementary contrasts. This can enable the extraction of more functional or structural information at any given anatomical location than what can be achieved with any single modality alone, which is a key advantage of MRI over other noninvasive imaging technologies. To achieve this, accurate alignment of these MRI data across the different contrasts acquired is critical. Head motion often occurs between scans, and rigid registration can be employed to align the datasets—by utilizing various available methods that can account for the different image contrasts—however this alignment is far more challenging in the presence of differential geometric distortion between these acquisitions. In many cases, these geometric distortions also can change with head position, further exacerbating error in alignment.

The most prevalent source of geometric distortion in MRI is inhomogeneity of the main magnetic field, B_0_, which for modern MRI scanners is predominantly induced by magnetic susceptibility differences within the volunteer’s head mainly found in regions near to air-tissue interfaces (e.g. at the sinuses, ear canals, and oral cavity). These local magnetic field variations cause field gradients that interfere with the image encoding gradients and thereby result in errors in image encoding such as voxel shift errors, which manifest as nonrigid geometric distortion (i.e., local expansion or compression) of the resulting image. For a given local B_0_ field offset, these image encoding errors scale with image encoding bandwidth, such that a lower encoding bandwidth results in a greater vulnerability to distortion and a larger voxel shift in the presence of field inhomogeneity.

Single-shot Echo Planar Imaging (EPI) is widely used for acquiring various MRI contrasts such as functional, diffusion and perfusion weighted imaging, and increasingly for anatomical imaging as well. EPI is well known to be vulnerable to these geometric distortions, which are primarily along the phase-encoding direction; in EPI, not only is this the encoding direction with the lowest bandwidth, but also in most applications of EPI the phase-encoding bandwidth is sufficiently low that distortions are clearly apparent, with maximum voxel displacements in the centimeter range. Phase encoding in EPI has relatively low encoding bandwidth (such as 30 Hz/pixel) compared to the relatively high frequency encoding bandwidths typically used in conventional (non-EPI) acquisitions (such as 600 Hz/pixel), thus geometric distortion is most often considered to be only in the phase-encoding direction in EPI, and for most non-EPI applications distortion in the frequency-encoding direction is negligible (Jezzard, 2012). While the distortion in the phase-encoding direction in EPI can be partly mitigated with parallel imaging acceleration (de Zwart et al., 2002; Griswold et al., 1999), which increases the effective phase-encoding bandwidth, the undersampling used in accelerated acquisitions brings with it a fixed penalty of decreased signal-to-noise ratio (SNR), and high acceleration factors can introduce severe artifacts even with modern receive coil arrays, therefore acceleration alone cannot fully remove the distortion in EPI. For this reason, EPI distortion must be addressed in post processing. Conventionally, post-hoc correction of the geometric distortion is performed based on an estimation of the voxel shifts induced by the B_0_ magnetic field inhomogeneity during the image acquisition, where the field offsets can be calculated from a separately acquired B_0_ map (Jezzard and Balaban, 1995). Another approach is based on the “reversed gradient” method that, instead of using an explicit B_0_ field map, infers the distortion using a pair of two EPI scans acquired with the phase-encoding gradients played out with opposite polarities, resulting in two images with equal magnitudes of distortion but in opposite directions, from which non-linear image registration can be used to simultaneously estimate and correct the distortion (Andersson et al., 2003; Bowtell et al., 1994; Chang and Fitzpatrick, 1992; Holland et al., 2010; Morgan et al., 2004).

While distortion in the phase-encoding and frequency-encoding directions have been thoroughly considered in previous work, the distortion in the slice-encoding direction for 2D slice-by-slice imaging has received some attention (Studholme et al., 2000), but is becoming more important as it may be increasingly present in modern MRI applications. For example, high-resolution (sub-millimeter) fMRI uses thin imaging slices which are usually acquired by maximizing slice-select gradient strength; however once the maximum strength is reached, a decrease of the slice-encoding gradient bandwidth is required to further narrow the range or excited frequencies, which could lead to the additional geometric distortions and “slice bending” in the slice direction. For conventional slice-selective pulses, slice thickness (TH) is determined by the slice-encoding gradient strength (G_z_) and slice-select RF pulse bandwidth (BW) such that TH=BW/γG_z_. While the value of G_z_ is usually dictated by the desired slice thickness, the RF pulse bandwidth is related to the pulse duration (*D*) and the time-bandwidth product (TBW), i.e., TBW ≡ *D* · BW. In sub-millimeter imaging, because G_z_ typically cannot be further increased, the slice thickness can be decreased by modifying the RF pulse design either by reducing TBW (which leads to degradation of the slice profile), or by increasing the pulse duration (which lowers SAR but increases the minimal echo time, TE). Thus, to decrease slice thickness, often one must resort to reducing the RF excitation pulse bandwidth. While stronger gradient coils help to achieve distortion-free thin slices, they bring a penalty of increased Peripheral Nerve Stimulation and potential eddy currents. High-resolution fMRI is commonly performed at ultra-high magnetic field strengths (≥7 T) where susceptibility-induced B_0_ field inhomogeneities are stronger than at conventional field strengths, suggesting that this imaging modality may be particularly vulnerable to slice distortion. In addition, in many applications a low slice-select RF pulse bandwidth may also be desirable to decrease power deposition (SAR). Slice distortion due to field inhomogeneity has previously been reported in MRI studies conducted using conventional field strengths as well (such as 1.5 T) (Sumanaweera et al., 1993) and in the presence of metal implants where magnetic susceptibility offsets are substantially higher than those between different tissue types (Hargreaves et al., 2011; Hopper et al., 2006; Lu et al., 2009).

To characterize this slice distortion and present a method for correcting it, here we use a novel acquisition that acquires pairs of images with reversed polarity of the slice-encoding gradient to create image pairs with equal slice distortions in opposing directions, and then demonstrate that typical 7T fMRI acquisitions do suffer from noticeable geometric distortion in the slice-encoding direction, which will result in alignment errors when registering these data to other datasets, e.g., to anatomical reference data typically acquired using 3D encoding. Our method is analogous to the reversed-gradient method used for removing distortion in the phase-encoding or frequency-encoding directions, and here it is applied in the slice-encoding direction. We demonstrate a correction of these slice distortions with our method, and evaluate it using both spin-echo (SE) or gradient-echo (GE) based data acquired with slice gradients and RF pulse (excitation and refocusing) bandwidths that are matched those used in the target fMRI protocol. Because we use, for correction, the popular *topup* method implemented in the FSL package (Andersson et al., 2003; Smith et al., 2004), we have named our method *overeasy* (Blazejewska et al., 2017). While we demonstrate our approach using high-resolution EPI data, it is applicable to any 2D multi-slice data suffering from geometric distortion in the slice direction, which becomes more pronounced at higher spatial resolutions and higher magnetic field strengths.

## Materials and methods

Seven healthy volunteers (4F/3M, 26±7 y.o.) participated in this study after providing written informed consent in accordance with our institution’s Human Research Committee; the study protocol was approved by the Institutional Review Board of Massachusetts General Hospital.

All data were acquired on a whole-body 7T scanner (MAGNETOM, Siemens Healthcare, Erlangen, Germany) equipped with a “short-z FOV” body gradient coil (SC72) and an inhouse-built 32-channel coil array for receive and a single-channel birdcage coil for transmit (Keil et al., 2010). During each experimental session multiple EPI scans were acquired following standard automatic B_0_ shimming.

The *overeasy* method used in this study consists of acquiring pairs of EPI data with reversed slice-select gradient polarity, with appropriate shifting of the RF pulse center frequency, which causes the voxel displacement direction to alternate between the two acquisitions, as illustrated in Figure 1. Note that the images were not acquired purely axially, and were tilted somewhat to be roughly aligned to the brain’s AC-PC axis, therefore the slice-select gradient was not along a single gradient axis thus the gradient was a combination of the physical G_x_, G_y_ and G_z_ gradients. Because the only parameter to change between this pair of images is the slice-select gradient polarity, the differential distortion between these images is in the direction perpendicular to the slice plane. Because these two images are acquired immediately after one another, we assume that the B_0_ inhomogeneity is identical between the two acquisitions, as is common for reversed-gradient methods. For each scanning session an additional B_0_ field map was also obtained for validation. See Table 1 for details of the scanning parameters.

**Figure 1.**
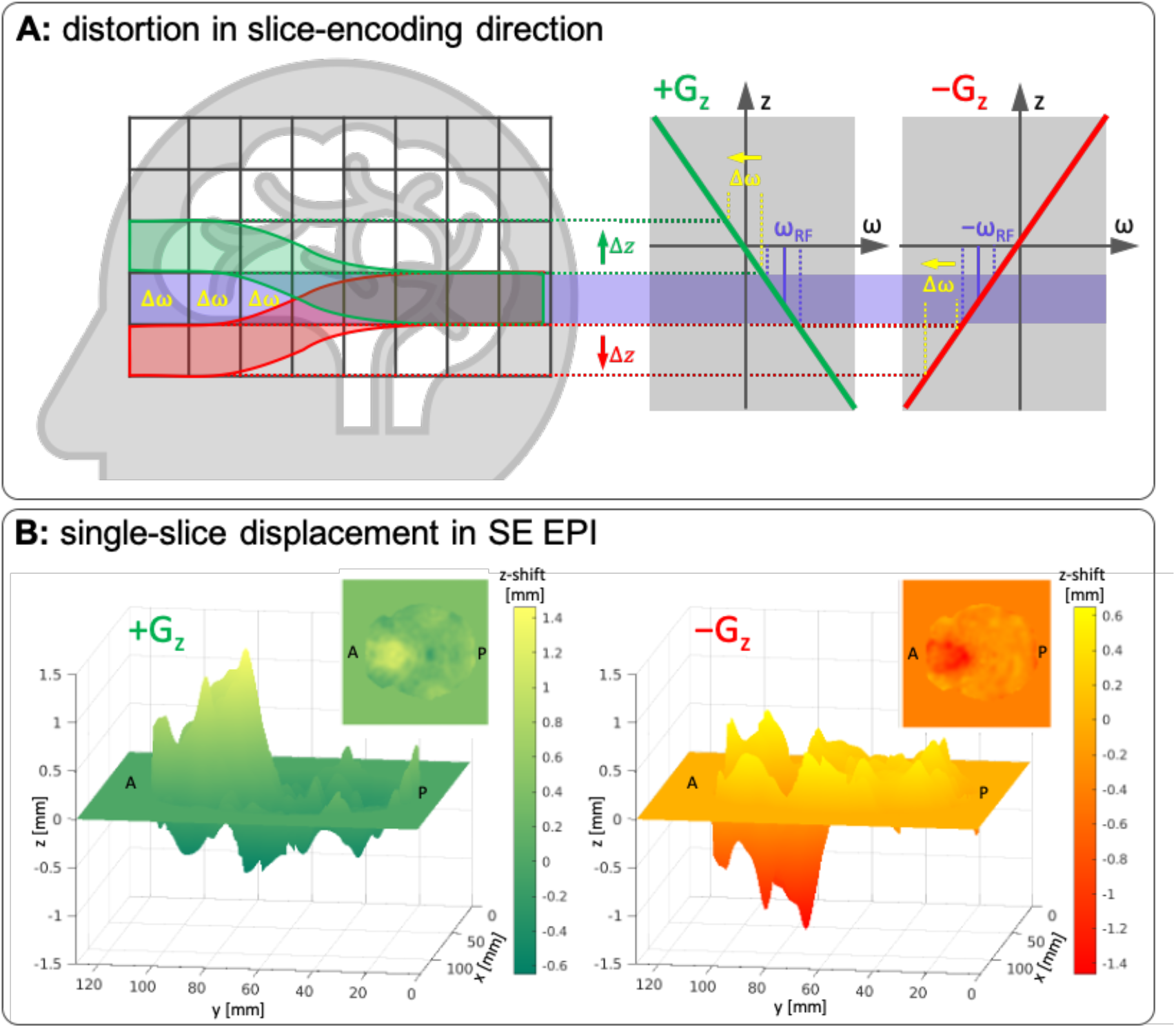
**(A)** Explanation of slice-encoding distortion due to local B_0_ field offsets and the reverse-gradient method applied to the slice-select direction. The slice-select gradient establishes a mapping between frequency and position along the slice-select direction (here, the z direction). A given slice location is excited by an RF pulse at the corresponding frequency. In the absence of local B_0_ field offsets, either a positive z gradient (+G_z_) or a negative z gradient (−G_z_) can be applied, and the desired slice location can be excited with an RF pulse using the appropriate center frequency, +ω_RF_ or −ω_RF_, respectively. In the presence of local B_0_ field offsets, the location of the excited slice will be shifted, and the direction of the shift will depend on the polarity of the slice-selective gradient. In this case, the direction of the shift will depend on whether the +G_z_ gradient or the −G_z_ gradient is used, and the magnitude of the shift will depend on the local B_0_ field offset Δω as well as the gradient amplitude and the RF pulse bandwidth. A desired slice location without the shift is marked with purple, while actual slice locations with the shift occurring in presence of +G_z_ and −G_z_ are marked with green and red, respectively. **(B)** Example of distortion-induced displacement (z-shift) of the voxels in the slice-encoding direction within a single axial slice, caused by local B_0_ field offsets, for images acquired with two opposite slice-encoding gradient polarities. In this example the shift is largest in the region at the front of the brain (near the frontal sinuses), and manifests as a positive z-shift when using the +G_z_ gradient and a negative z-shift when using the −G_z_ gradient.

**Table 1.** Summary of the EPI protocol parameters: TBW = time-bandwidth product in the slice-encoding direction, D = duration of a slice-encoding pulse, Gz = polarities of slice encoding, Gy = directions of phase-encoding gradient (where ‘ap’ stands for anterior-to-posterior and ‘pa’ stands for posterior-to-anterior), TH = slice thickness, RES = nominal in-plane voxel size, TR = repetition time, TE = echo time(s), FA = flip angle, ESP = nominal EPI echo spacing, R = in-plane acceleration factor, TA = time of acquisition, SL = number of slices, FOV = field of view. The right section of the table marks which data were acquired for each of the subjects.

| sequence | TBW | D [ms] | Gz | Gy | TH [mm] | RES [mm] | TR [ms] | TE(s) [ms] | FA [o] | ESP [ms] | R | TA | SL | FOV [mm] | ss02 | ss03 | ss04 | ss05 | ss06 | sg02 | sg03 |
| --- | --- | --- | --- | --- | --- | --- | --- | --- | --- | --- | --- | --- | --- | --- | --- | --- | --- | --- | --- | --- | --- |
| EPI GRE | 2.6 | 6/8/10 | pos/neg | ap | 1.0 | 1.5 | 5940 | 20 | 75 | 0.66 | 4 | 1:05 | 128 | 192 x 192 | ✓ | ✓ | ✓ | ✓ | ✓ | ✓ | ✓ |
| EPI GRE | 5.2/7.2 | 10 | pos/neg | ap | 1.0 | 1.5 | 5940 | 20 | 75 | 0.66 | 4 | 1:05 | 128 | 192 x 192 |  |  |  | ✓ | ✓ | ✓ | ✓ |
| EPI SE | 2.6 | 6/8/10 | pos/neg | ap | 1.0 | 1.5 | 15970 | 50 | 90 | 0.66 | 4 | 2:56 | 128 | 192 x 192 | ✓ | ✓ | ✓ | ✓ | ✓ |  |  |
| EPI SE | 2.6 | 10 | pos/neg | ap/pa | 1.0 | 2.0 | 15970 | 50 | 90 | 0.66 | 2 | 2:08 | 128 | 192 x 192 | ✓ | ✓ | ✓ | ✓ | ✓ |  |  |
| B0 map | -- | -- | -- | ap | 2.0 | 2.0 | 570 | 3.0/4.02 | 46 | -- | -- | 1:52 | 128 | 192 x 192 | ✓ | ✓ | ✓ | ✓ | ✓ | ✓ | ✓ |

Both GE and SE variants of the *overeasy* acquisition were tested, each with similar EPI readouts and slice bandwidths. The spin-echo EPI may be advantageous over gradient-echo EPI at 7T because of the potential for through-slice dephasing in the gradient-echo, which could cause signal loss exactly in those regions that suffer the most from distortion.

Each EPI scan consisted of 5 repetitions (for noise averaging) and the two slice-encoding polarity scans were acquired consecutively. Both the SE and GE EPI data were acquired near-axially (see above) at 1.5 mm in-plane resolution and a slice thickness of 1 mm using TBW=2.6. To demonstrate the effect of slice bandwidth on the slice distortion, we acquired multiple runs with RF pulse durations *D*=6, 8 and 10 ms (which corresponds to BW=433, 325 and 260 Hz, respectively), and for a subset of four subjects the GE data with TBW=5.2 and 7.2 and *D*=10 ms (BW=520 and 720 Hz, respectively).

To test whether the *overeasy* method could be used to simultaneously estimate slice-encoding distortion and phase-encoding distortion, for a subset of 5 subjects we also acquired SE EPI data where we alternated both the slice-select gradient polarity and the phase-encoding gradient polarity. These data were acquired with 2 mm in-plane resolution and 1-mm slice thickness using TBP=2.6 and *D*=10 ms, with alternating phase-encoding directions: anterior-to-posterior and posterior-to-anterior (four datasets in total).

To allow us to directly compare the distortion-correction results across all acquired slice-encoding bandwidths and between the SE and GE EPI data, for each subject all datasets acquired with positive slice-encoding gradients were co-registered together and then all datasets acquired with negative slice-encoding gradients were co-registered together using rigid (i.e., 6 degrees of freedom) registration calculated using the FLIRT tool of the FSL software package (Jenkinson et al., 2002; Jenkinson and Smith, 2001). The voxel shift deformation fields were calculated for each reversed slice-encoding volume pair using the *topup* reversed-gradient method as implemented in FSL (Andersson et al., 2003; Smith et al., 2004) from SE and GE data separately. While *topup* is mainly intended for correcting distortion along the phase-encoding direction, it is possible to use this software to correct distortion in any encoding direction, therefore no modifications to the software were needed. The *topup* software performs simultaneous motion correction of the input dataset to correct for potential distortion-independent shifts between volumes acquired with reversed slice-encoding gradient polarities; this effectively provides a co-registration across positive and negative slice-encoding gradient polarity volume pairs, ignoring any potential motion in the distorted direction. Two distortion correction approaches were evaluated and applied to GE EPI data: (1) using the voxel shifts in the slice direction estimated from the SE EPI *overeasy* pairs, consistent with the typical use case of the *topup* method, and (2) using voxel shifts in the slice direction estimated from the GE EPI data themselves, to test whether any signal loss present due to through-slice dephasing would bias the results.

Slice-distortion can also be estimated using a conventional B_0_ field map, provided that the slice-encoding bandwidth and gradient direction are known, and this distortion estimate should be equivalent to that given by *overeasy*. Therefore, to validate the *overeasy*-based correction, voxel shift maps based on separately acquired B_0_ field map scans resampled to the EPI space were calculated, then distorted along the EPI phase-encoding direction so that they more accurately matched the distorted EPI data. The resulting voxel shift maps were estimated for different TBW and *D* combinations and compared pairwise with those estimated from the corresponding *overeasy* data at a voxel-by-voxel basis, and the consistency between these two estimates was evaluated by calculating the correlation coefficient between the B_0_-fieldmap-based and the *overeasy*-based estimates.

Finally, a two-step *topup* approach was used to correct both slice-encoding and phase-encoding distortion an additional SE EPI dataset acquired with two pairs of reversed gradient directions—i.e., the SE EPI data were acquired four times with both positive and negative phase-encoding and slice-encoding directions. For correction, the phase-encoding distortion correction was performed first using the two pairs of data, after which the slice-encoding distortion correction was performed.

## Results

We found clear geometric distortion to occur in the slice-encoding direction in all GE and SE EPI datasets acquired with the proposed *overeasy* method which allows for reversing slice-select gradient polarity, as illustrated in Figure 1. Example pairs of images with alternating slice-encoding directions are shown in Figure 2, where distortions up to several slice-thicknesses are observable in regions of strong B_0_ field inhomogeneity. The regions near the frontal sinus where distortion is the most severe were magnified to visualize the magnitude of voxel shifts; to aid in the visual inspection of these displacements, a horizontal red line—placed in the same location in scanner coordinates across the two images of the *overeasy* pairs—is included as a reference. This simple reversal of the slice-encoding gradient polarities added into the pulse sequences provides a concrete demonstration of the slice-distortion effect. We then corrected the slice-select direction distortion in the *overeasy* pairs of images by applying *topup* method, where either SE or GE EPI reference data of opposite slice-encoding gradient polarities were used to derive a deformation field (Figure 3).

**Figure 2.**
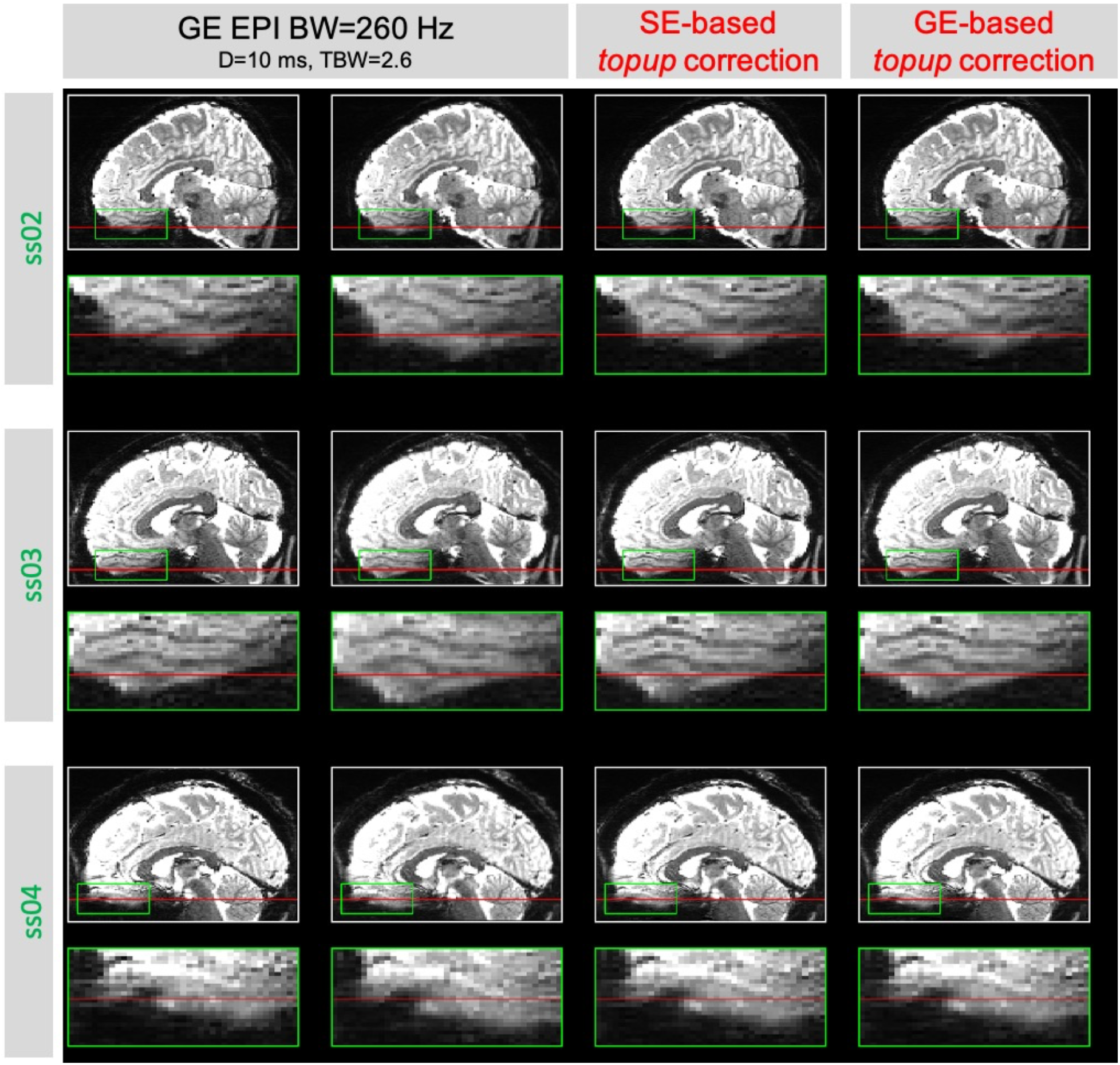
Example sagittal reformats of GE EPI datasets from three different subjects (ss02, ss03 and ss04), acquired with 1-mm slice thickness, 1.5-mm in-plane resolution, positive and negative slice-encoding gradient direction, slice-encoding pulse duration *D*=10 ms and TBW=2.6, which corresponds to BW=260 Hz (left). The positive (+G_z_) and negative (–G_z_) slice encoding directions are indicated by red arrows. Corresponding slices distortion-corrected using SE-based (middle) and GE-based (right) *data*. The green box represents the regions shown in the magnified inserts, which show distortion in the slice encoding direction in the frontal sinus region, and the red horizontal line is positioned at the same location in all images as a reference to help visualize the amount of displacement in the image data. The corresponding images for all remaining subjects and BW values tested are presented in Supplementary Figure 1.

**Figure 3.**
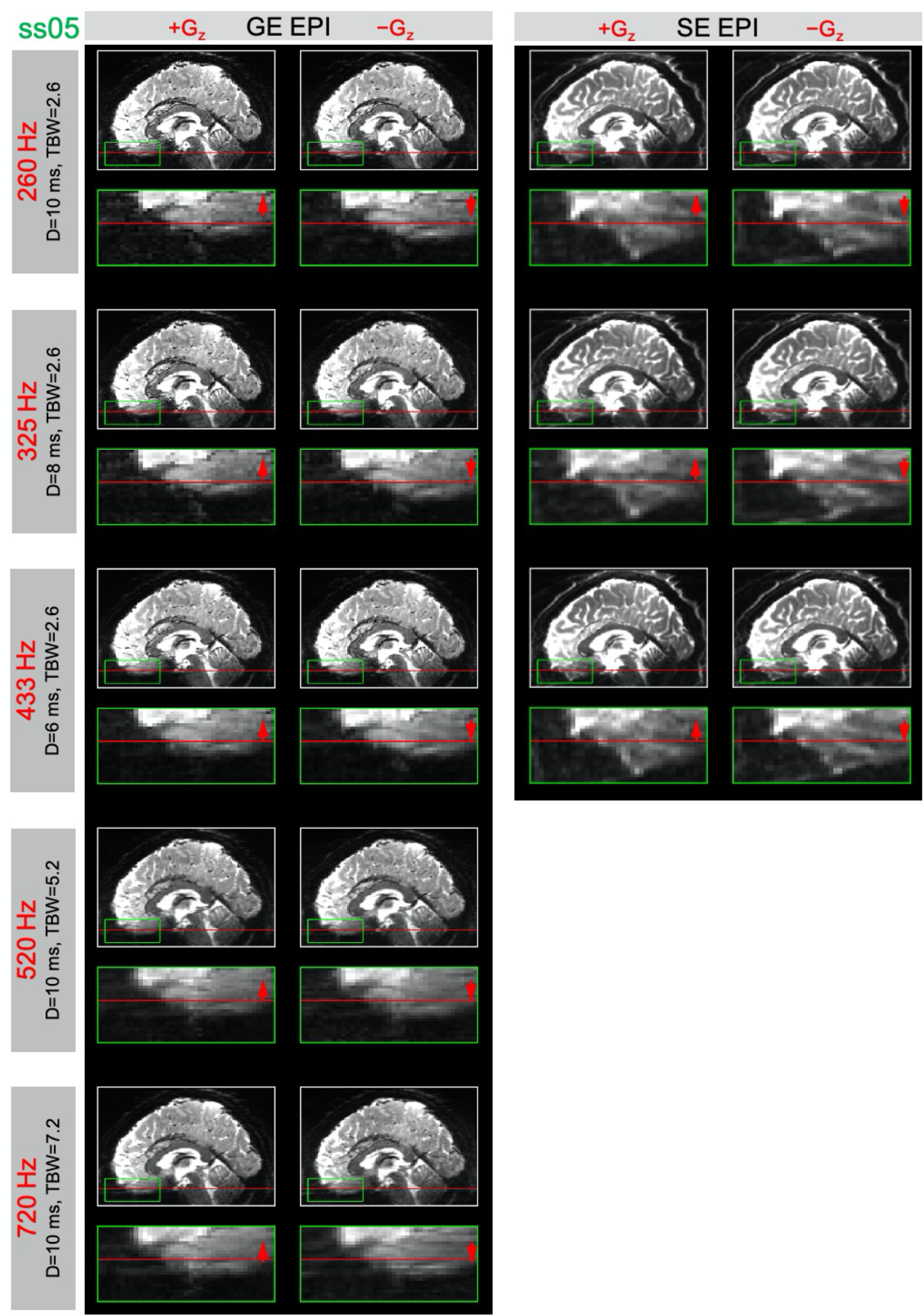
Example sagittal reformats of gradient-echo (GE, left) and spin-echo (SE, right) EPI datasets from a representative subject (ss05), with 1-mm slice thickness and 1.5-mm in-plane resolution. Positive (+G_z_, left) and negative (–G_z_, right) slice-encoding direction (indicated by red arrows). The rows are ordered according to increasing slice-encoding BW values achieved by changing either the slice-select/slice-refocus pulse duration (*D*) or timebandwidth product (TBW); see text for details. The horizontal red lines are included in the same position in each pair of images to help visualize shift of voxels due to distortion; magnified inserts correspond to a region indicated by the green box where distortion is pronounced. A summary of the data acquired for all remaining subjects is presented in Supplementary Figure 2.

Figure 2 presents representative example sagittal reformats of the original distorted GE EPI data acquired with TBW=2.6 and *D*=10 ms (BW=260 Hz) with positive and negative gradient polarities (marked with red arrows at the images) for three different subjects, together with the corrected data based on either the deformation estimated from the SE EPI data or from the GE EPI data themselves. An overview of the distorted and corresponding corrected data for all remaining subjects is presented in Supplementary Figure 1.

As expected, the extent of distortion in the slice-encoding direction was the most pronounced in the images acquired with the lowest slice-encoding gradient bandwidth of 260 Hz, which corresponded to TBW of 2.6 and *D* of 10 ms, and gradually decreased with increasing bandwidth values. Figure 3 presents a comparison of the example sagittal reformats from GE and SE EPI datasets acquired for one subject with different BW values ranging from 260 to 720 Hz, and both positive and negative slice-encoding gradient polarity. A summary of the examples for all the datasets and the corresponding distortion corrected slices from all remaining subjects can be found in Supplementary Figure 2. Nearly identical results were obtained from our SE and GE EPI data as shown in Figure 4 which summarized the correlations between the voxel shift maps resulting from both *topup* approaches and the voxel shift map derived from the field map scan. The correlation values are similar for SE- and GE-*topup*, suggesting that SE EPI data may not be needed when using GE data with such thin slices, even at 7T.

**Figure 4.**
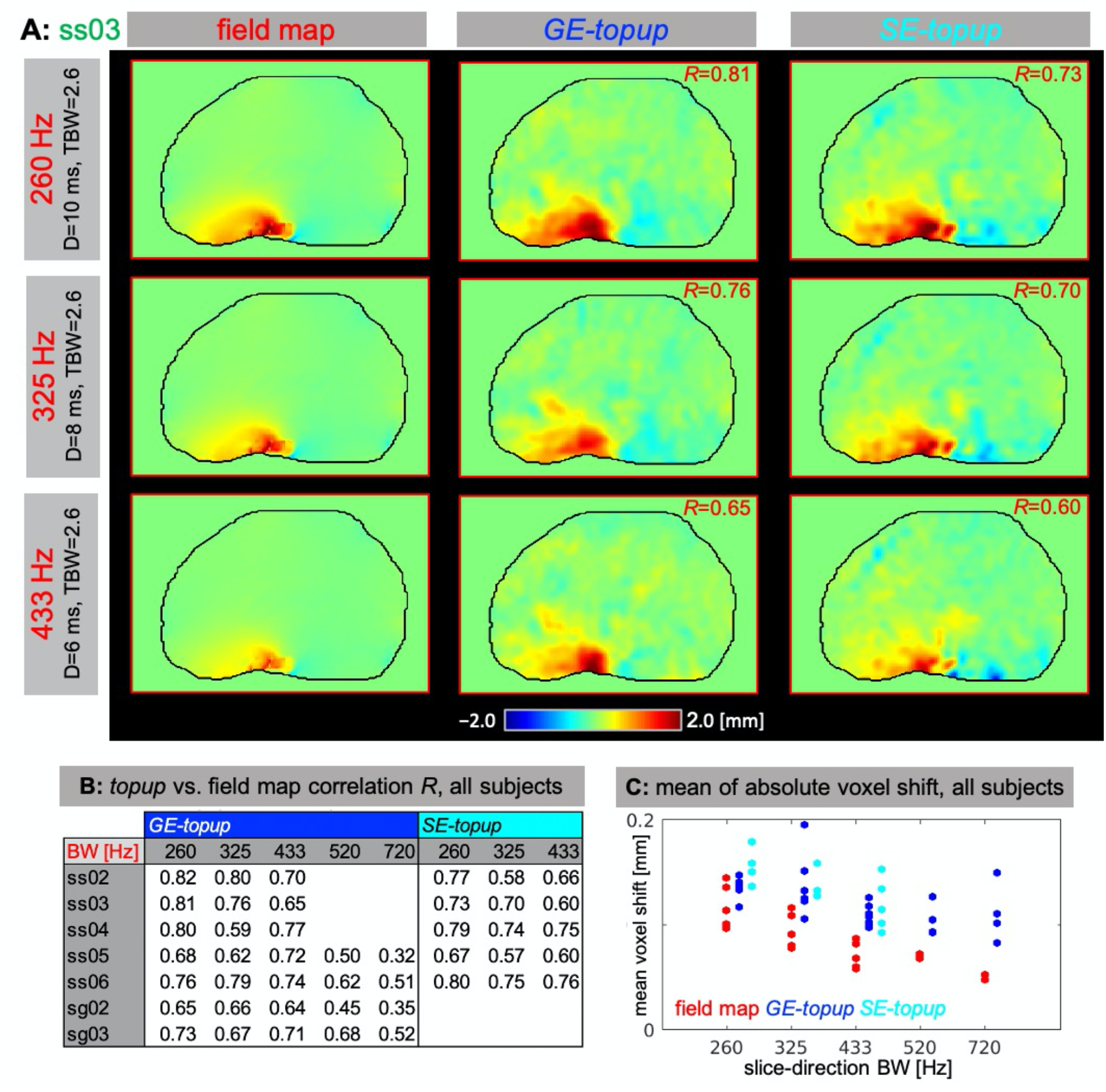
(**A)** Example sagittal reform ats of voxel shift maps in the slice-encoding direction calculated for a representative subject (ss03) using the reverse-gradient-based (i.e., *topup*) (left) and conventional B_0_ field map-based approach (middle) accompanied by the correlation coefficient calculated for each pair within the brain region (right). **(B)** Summary of the correlation coefficients calculated for pairs of the voxel shift maps across all subjects and BW values. **(C)** Comparison of *topup* and field map-based mean absolute voxel shift values for all subjects and BWs, with the blue points representing the voxel shift estimated from *topup* and the cyan points representing the corresponding voxel shift estimated from the B_0_ field map.

All voxel shift maps generated using *topup* and calculated from the separately acquired B_0_ field map scans demonstrated similar spatial distribution of the shift values, reaching ±2 mm which corresponds to two times the slice thickness, as can be seen in Figure 4A for a representative subject. The correlation coefficients calculated between *topup* and field map-based maps within the brain mask were highest for the lowest BW values (the most distorted datasets) reaching 0.58–0.82 (Figures 4A & B) and the correlation coefficient decreased with the increasing BW. The lower correlation coefficient seen for the data with the lowest level of distortion is likely due to the smaller displacements estimated in the presence of noise (i.e., lower “displacement-to-noise ratio”). The mean values of absolute voxel shifts calculated within the brain were also similar for both approaches, with *topup* estimations being slightly higher overall and *SE-topup* estimations being slightly higher than *GE-topup*. (Figure 4C). Images acquired with lower slice-gradient BW demonstrated larger mean absolute voxel shift values (Figure 4C) over larger spatial area of the brain (Figure 4A) with the decreasing trend for increasing BWs, as expected.

We also demonstrated a two-step approach using *topup* to correct distortion in phase-encoding and slice-select direction applied to four SE EPI volumes acquired separately with A→P and P→A phase-encoding gradient direction, and positive and negative slice-encoding gradient direction (Figure 5).

**Figure 5.**
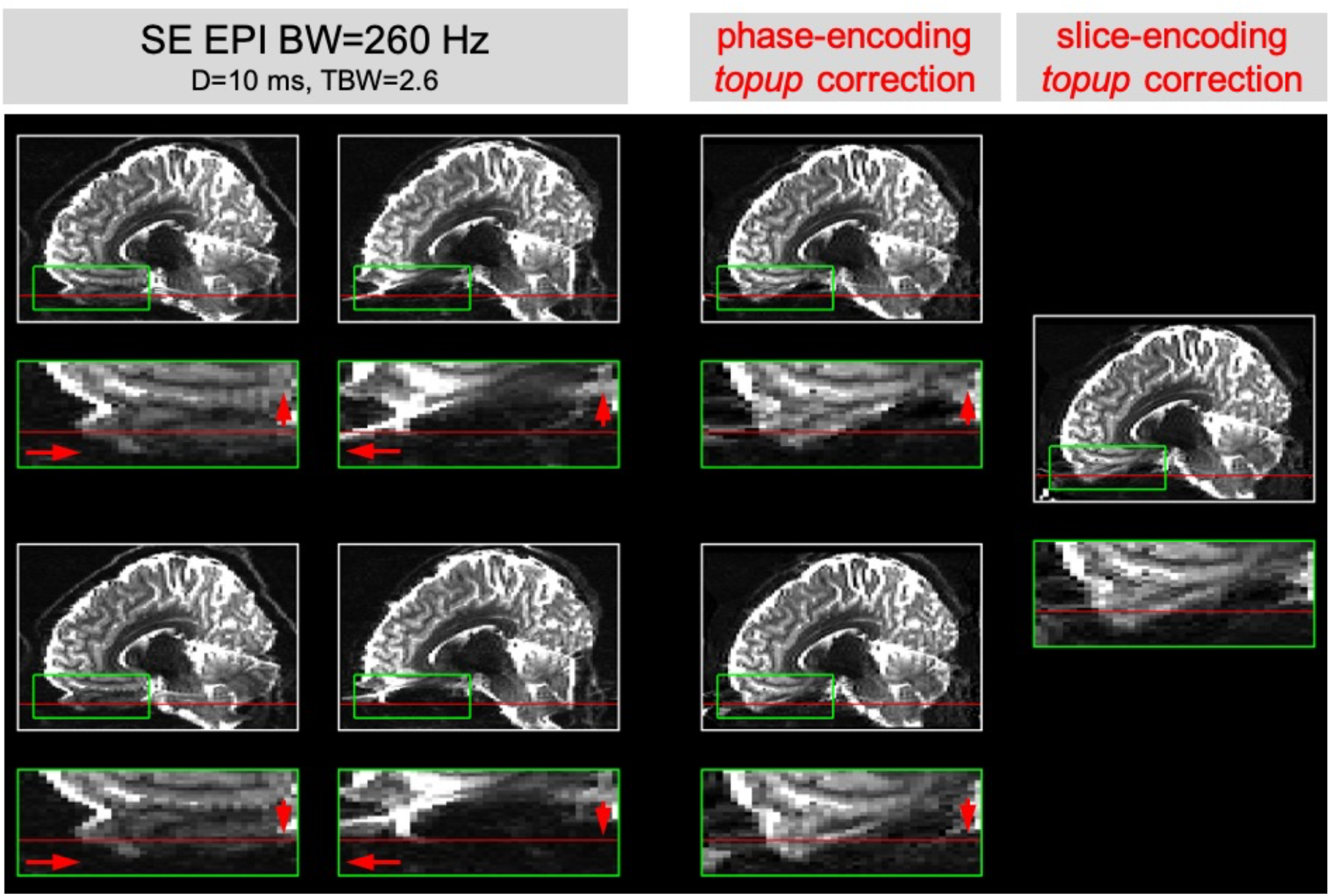
Example sagittal reformats of the SE EPI dataset, acquired axially with 1-mm slice thickness, 2.0 mm in-plane resolution, A→P and P→A phase-encoding gradient direction (first and second column, respectively), positive and negative slice-encoding gradient direction (top and bottom row, respectively), slice-encoding pulse duration D=10 ms and TBW=2.6, which corresponds to BW=260 Hz; positive (+G_z_) and negative (−G_z_) slice-encoding direction, as well as A→P and P→A phase-encoding direction are indicated by red arrows; corresponding data distortion-corrected using *topup* method in phase-encoding direction (third column) and slice-encoding direction (fourth column); magnified inserts show distortion in the frontal sinus region with red horizontal line positioned in the same location in all panels.

## Discussion

In this work we presented a simple method for quantifying distortion in the slice-select direction, and demonstrated that modern high-resolution (i.e., thin-sliced) 7T fMRI data contain geometric distortion in the slice direction, with a voxel shifts that may exceed the slice thickness. We were able to measure these distortions from either SE or GE-based acquisitions, indicating that for these thin-slice protocols through-slice dephasing does not prevent accurate estimations from GE-based data, although whether this finding generalizes to other GE-based protocols will depend on several factors including slice thickness, TE, slice prescription, brain region, and B_0_ shim quality. While these slice-encoding distortions are smaller in magnitude than the well-known phase-encoding distortions, for accurate alignment of data across different MRI data, such as alignment of 2D EPI data to 3D anatomical reference data, slice distortion must be addressed to ensure accuracy of the alignment.

While slice distortion can be readily estimated using conventional B_0_ field map methods, provided the slice-select bandwidth and gradient direction are known, here we used a simple approach to demonstrate in a concrete way the level of slice-distortion that does not require any knowledge of the RF pulses. Our method, based on pairs of EPI data, is faster than conventional methods that acquire an explicit B_0_ field map, which generally require multiple minutes, making them less practical—especially in cases where estimating distortion multiple times during a session is desirable, which may be the case in subjects prone to motion. Because magnetic-susceptibility-induced B_0_ inhomogeneity changes with head position, and, in particular, with head orientation relative to the B_0_ axis, updating the B_0_ inhomogeneity and the associated image distortion periodically during the imaging session can help ensure consistent accuracy of the correction throughout the session. EPI-based field maps are another possibility and have some advantages over B_0_ field maps acquired with conventional encoding (Hutton et al., 2002), provided that the distortions are not too severe. These have also been extended to estimating dynamic changes in distortion during fMRI runs (Lamberton et al., 2007). Still, an advantage of acquiring pairs of reverse-gradient images is that it provides a means to validate distortion correction based on conventional B_0_ field maps: if the B_0_ field map yields an accurate estimate of the image distortion, applying the B_0_-fieldmap-based correction to each image of the reversed-gradient pair should result in two identical corrected images. Also, it is possible to estimate slice-encoding distortion present in conventional (i.e., non-EPI) image encoding methods as well using the *overeasy* method, time permitting, if the slice-select RF pulse bandwidth can be set to be identical to that of the target acquisition.

Another important advantage of reverse-gradient methods for estimating slice-encoding distortion is that they can also help remove distortions due to eddy currents. Since the slice-select gradient amplitude can be larger than other gradient blips, it can be a source of eddy currents that induce B_0_ inhomogeneity in the image and therefore distortion (Varadarajan et al., 2021). Reversing the slice-encoding gradient will reverse the eddy currents and therefore both eddy current and susceptibility distortions can be corrected. This is challenging to achieve with conventional B_0_ field maps, unless again the B_0_ field map uses the same slice-select gradient as the target acquisition so that both the B_0_ field map and the target acquisition experience the same eddy currents from the slice-select gradient, which would allow the B_0_ field map to measure the same B_0_ inhomogeneity (due to both eddy currents and susceptibility gradients) that is present during the target acquisition.

When B_0_ inhomogeneity becomes severe, the distortions along the phase-encoding direction can grow sufficiently large as to induce “singularities” in the EPI data where the compressive distortion results in what is known as “voxel pile-up” artifacts. These artifacts cannot be corrected when using a single phase-encoding polarity, but they can be resolved when reversed-gradient pairs are available. In principle it is possible for “slice pile-up” to occur, although given that the slice-encoding bandwidth is considerably higher (20× and more) than typical phase-encoding bandwidths this is unlikely. In more moderate cases, B_0_ inhomogeneity will cause a spatially-varying voxel size in the phase-encoding direction, due to the geometric expansion and compression of the image (Wang et al., 2021). While B_0_ inhomogeneity is often viewed as causing mainly a shift in the slice position, if the B_0_ field changes *within the slice* along the slice direction it can similarly cause an expansion or compression. However, given that relatively low slice-encoding bandwidths are mainly used to achieve thin slices, and the B_0_ field is generally smoothly varying in space, it is also unlikely that the B_0_ field will change considerably at the spatial scale of these thin slices. Future work will consider the effects of B_0_ inhomogeneity on the true resolution of the image in the slice direction.

A final practical consideration is on how to apply this distortion correction to the data. While here the image pairs were corrected directly, using a nonlinear warping followed by an interpolation, in practice, especially in high-resolution applications, it is preferable to mathematically compose all transformations of the data to minimize the number of interpolation steps (Glasser et al., 2013; Polimeni et al., 2018). Because the distortion in the slice-encoding direction will typically be far smaller than that of the phase-encoding direction, most of the displacements in the slice-encoding direction will likely be sub-voxel shifts, which will induce blurring when interpolation is performed. Higher-order interpolation, combined with image grid upsampling, can also help mitigate resolution losses due to interpolation.

Here we focused on demonstrating the distortion in the slice-encoding direction by reversing the gradient polarity of the slice-select gradient, and demonstrated that the bandwidths used in modern fMRI acquisitions do lead to perhaps unexpectedly large levels of distortion the slice direction. This approach can be combined with existing reverse-gradient methods that reverse the phase-encoding direction, as we demonstrated above (see Figure 5). While here we acquired four images, but reversing each gradient direction, in principle only one pair is needed, e.g., a positive-slice-encode and positive-phase-encode image followed by a negative-slice-encode and negative-phase-encode image, to save time. In this case, the differential distortion between the two images is no longer constrained to be only along the slice-encoding direction or along the phase-encoding direction, rather it would be at an angle partway between these two directions, where this angle would depend on the ratio of the bandwidths. For example, if the phase-encoding direction was in *y*, the slice-encoding in *z*, and the phase-encoding bandwidth were 5 times larger than the slice-encoding bandwidth, the differential distortion between the two images of the pair would be along a line about tan^−1^(1/5) = 11° off of the *y* axis.

## Conclusions

Distortion in the slice-encoding direction up to several millimeters was detected in data acquired at 7T using a standard high-resolution fMRI protocol. Reducing bandwidth in the slice-encoding direction allows for acquiring thin slices but introduces vulnerability to geometric distortion due to B_0_ inhomogeneity. While this distortion is lower than that found in the phase-encoding direction, it is large enough to reduce accuracy in the alignment of data acquired using different protocols. Both GE and SE EPI acquisitions were used to acquire reversed-gradient data by alternating the polarity of the slice-encoding gradient along with the RF pulse frequency offset. These pairs of data allowed distortion correction using a conventional reverse-gradient method, implemented in *topup*. This slice-encoding gradient reversal can also be used as a means to validate geometric distortion correction based on acquired B_0_ field maps. The proposed reversed-gradient-based distortion correction methods, termed *overeasy*, is beneficial in that it reduces the scan time needed to estimate distortion, just as existing EPI-based reversed-gradient methods do for phase-encoding distortion correction, and helps ensure that the relevant distortions from both susceptibility effects and eddy currents are estimated properly.

## Acknowledgements

Special thanks to Ned Ohringer and Kyle Droppa for help with subject recruitment and to Azma Mareyam and Dr. Jason Stockman for hardware support. This research has been supported by NIH NIBIB (P41-EB015896, P41-EB030006 and R01-EB019437), by the *BRAIN Initiative* (NIMH R01-MH111419 and K99-MH120054), and by the Athinoula A. Martinos Center for Biomedical Imaging, and made possible by NIH Shared Instrumentation Grants S10-RR023401, S10-RR023043 and S10-RR020948.

## References

Andersson, J.L.R., Skare, S., Ashburner, J., 2003. How to correct susceptibility distortions in spin-echo echo-planar images: application to diffusion tensor imaging. Neuroimage 20, 870–88. https://doi.org/10.1016/S1053-8119(03)00336-7

Blazejewska, A.I., Witzel, T., Wald, L.L., Polimeni, J.R., 2017. Correction of EPI geometric distortion in slice direction using reversed slice-select gradients and topup. Proc. Int. Soc. Magn. Reson. Med. 25, 1650.

Bowtell, R.W., McIntyre, D.J.O., Commandre, M.-J., Glover, P.M., Mansfield, P., 1994. Correction of Geometric Distortion in Echo Planar Images, in: ISMRM. p. 441.

Chang, H., Fitzpatrick, J.M., 1992. A Technique for Accurate Magnetic Resonance Imaging in the Presence of Field Inhomogeneities. IEEE Trans. Med. Imaging 11, 319–329. https://doi.org/10.1109/42.158935

de Zwart, J.A., Van Gelderen, P., Kellman, P., Duyn, J.H., 2002. Application of sensitivity-encoded echo-planar imaging for blood oxygen level-dependent functional brain imaging. Magn. Reson. Med. 48, 1011–1020. https://doi.org/10.1002/mrm.10303

Glasser, M.F., Sotiropoulos, S.N., Wilson, J.A., Coalson, T.S., Fischl, B., Andersson, J.L., Xu, J., Jbabdi, S., Webster, M., Polimeni, J.R., Van Essen, D.C., Jenkinson, M., 2013. The minimal preprocessing pipelines for the Human Connectome Project. Neuroimage 80, 105–124. https://doi.org/10.1016/j.neuroimage.2013.04.127

Griswold, M.A., Jakob, P.M., Chen, Q., Goldfarb, J.W., Manning, W.J., Edelman, R.R., Sodickson, D.K., 1999. Resolution enhancement in single-shot imaging using simultaneous acquisition of spatial harmonics (SMASH). Magn. Reson. Med. 41, 1236–1245. https://doi.org/10.1002/(SICI)1522-2594(199906)41:6<1236::AID-MRM21>3.0.CO;2-T

Hargreaves, B.A., Worters, P.W., Pauly, K.B., Pauly, J.M., Koch, K.M., Gold, G.E., 2011. Metal-induced artifacts in MRI. Am. J. Roentgenol. 197, 547–555. https://doi.org/10.2214/AJR.11.7364

Holland, D., Kuperman, J.M., Dale, A.M., 2010. Efficient Correction of Inhomogeneous Static Magnetic Field-Induced Distortion in Echo Planar Imaging. 2NeuroImage 50, 1–18. https://doi.org/10.1016/j.neuroimage.2009.11.044.Efficient

Hopper, T.A.J., Vasilić, B., Pope, J.M., Jones, C.E., Epstein, C.L., Song, H.K., Wehrli, F.W., 2006. Experimental and computational analyses of the effects of slice distortion from a metallic sphere in an MRI phantom. Magn. Reson. Imaging 24, 1077–1085. https://doi.org/10.1016/j.mri.2006.04.019

Hutton, C., Bork, A., Josephs, O., Deichmann, R., Ashburner, J., Turner, R., 2002. Image distortion correction in fMRI: A quantitative evaluation. Neuroimage 16, 217–40. https://doi.org/10.1006/nimg.2001.1054

Jenkinson, M., Bannister, P., Brady, M., Smith, S., 2002. Improved optimization for the robust and accurate linear registration and motion correction of brain images. Neuroimage 17, 825–41.

Jenkinson, M., Smith, S., 2001. A global optimisation method for robust affine registration of brain images. Med. Image Anal. 5, 143–156.

Jezzard and Balaban, 1995. Correction for Geometric Distortion in Echo Planar Images from B0 Field Variations. Magn. Reson. Med. 34, 65–73.

Jezzard, P., 2012. Correction of geometric distortion in fMRI data. Neuroimage 62, 648–51. https://doi.org/10.1016/j.neuroimage.2011.09.010

Keil, B., Triantafyllou, C., Hamm, M., Wald, L.L., 2010. Design optimization of a 32-channel head coil at 7T, in: Intl Soc Mag Reson Med. p. 1493.

Lamberton, F., Delcroix, N., Grenier, D., Mazoyer, B., Joliot, M., 2007. A new EPI-based dynamic field mapping method: application to retrospective geometrical distortion corrections. JMRI 26, 747–55. https://doi.org/10.1002/jmri.21039

Lu, W., Pauly, K.B., Gold, G.E., Pauly, J.M., Hargreaves, B.A., 2009. SEMAC: Slice Encoding for Metal Artifact Correction in MRI. Magn. Reson. Med. 62, 66–76. https://doi.org/10.1002/mrm.21967.SEMAC

Morgan, P.S., Bowtell, R.W., McIntyre, D.J.O., Worthington, B.S., 2004. Correction of spatial distortion in EPI due to inhomogeneous static magnetic fields using the reversed gradient method. JMRI 19, 499–507. https://doi.org/10.1002/jmri.20032

Polimeni, J.R., Renvall, V., Zaretskaya, N., Fischl, B., 2018. Analysis strategies for high-resolution UHF-fMRI data. Neuroimage 168, 296–320. https://doi.org/10.1016/j.neuroimage.2017.04.053

Smith, S.M., Jenkinson, M., Woolrich, M.W., Beckmann, C., Behrens, T.E.J., Johansen-Berg, H., Bannister, P.R., De Luca, M., Drobnjak, I., Flitney, D.E., Niazy, R.K., Saunders, J., Vickers, J., Zhang, Y., De Stefano, N., Brady, J.M., Matthews, P.M., 2004. Advances in functional and structural MR image analysis and implementation as FSL. Neuroimage 23 Suppl 1, S208–19.

Studholme, C., Constable, R.T., Duncan, J.S., 2000. Accurate alignment of functional EPI data to anatomical MRI using a physics-based distortion model. IEEE Trans. Med. Imaging 19, 1115–27. https://doi.org/10.1109/42.896788

Sumanaweera, T.S., Glover, G.H., Sumanaweera, T.O., Adler, J.R., 1993. MR susceptibility misregistration correction. IEEE Trans. Med. Imaging 12, 251–259. https://doi.org/10.1109/42.232253

Varadarajan, D., Balasubramanian, M., Park, D.J., Witzel, T., Stockmann, J.P., Polimeni, J.R., 2021. Characterizing the acquisition protocol dependencies of B0 field mapping and the effects of eddy currents and spoiling. Proc. Int. Soc. Magn. Reson. Med. 29, 3552.

Wang, J., Nasr, S., Roe, A.W., Polimeni, J.R., 2021. Evaluation of spatial blur induced by preprocessing and distortion in UHF fMRI data. Proc Intl Soc Mag Reson Med 29, 3126.

