## Supplementary figures for "Slice-direction geometric distortion evaluation and correction with reversed slice-select gradient acquisitions"

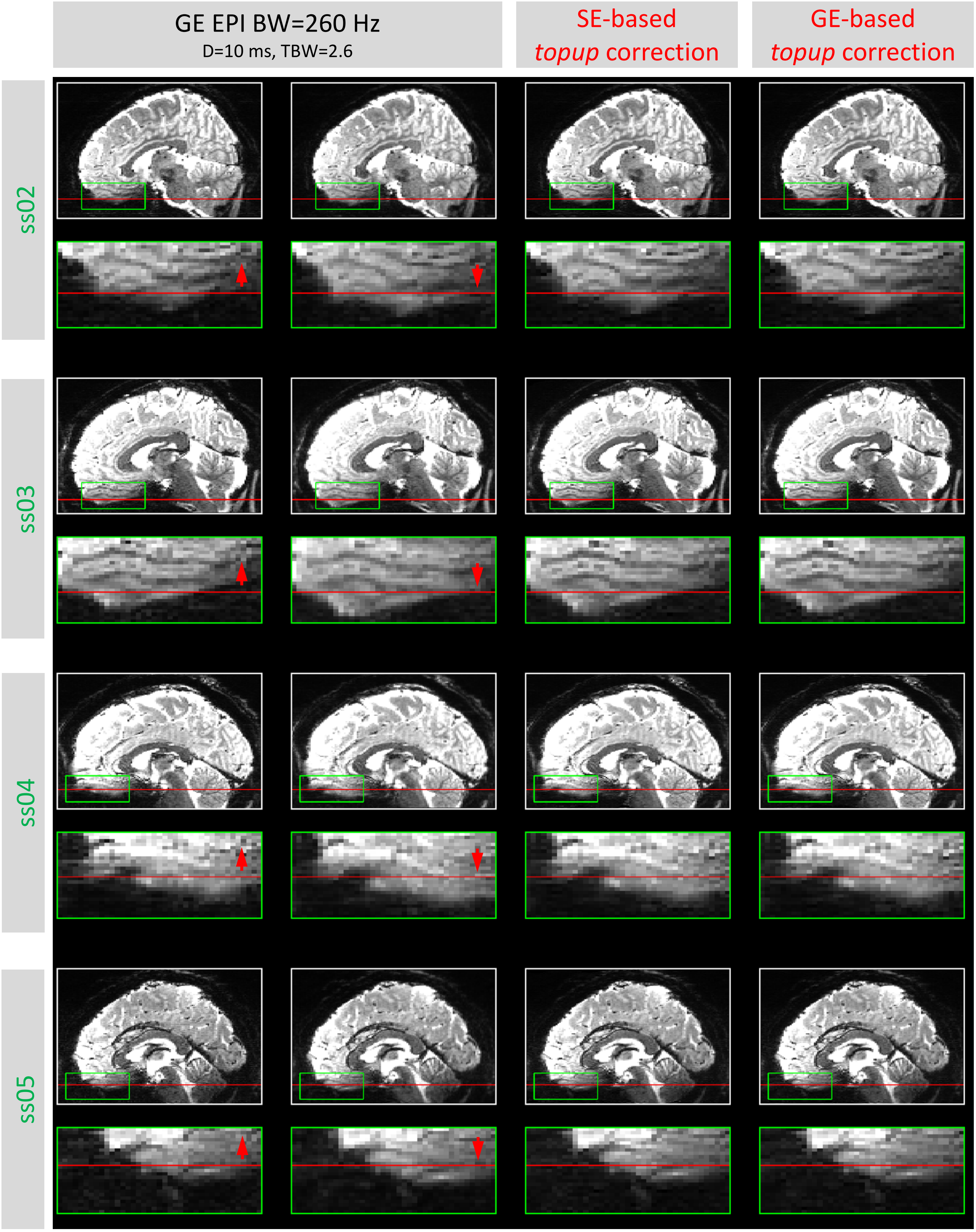


**Supplementary Figure 1.** Example sagittal reformats of gradient-echo EPI scans from all subjects, with positive and negative slice encoding direction and different combinations of slice-encoding pulse duration and TBW (two left columns), with corresponding *topup* distortion-corrected datasets (two right columns), as shown in Figure 2. Inserts show magnification of region within green box to better visualize the slice-encoding distortion in a region of large $B_0$ field offsets.

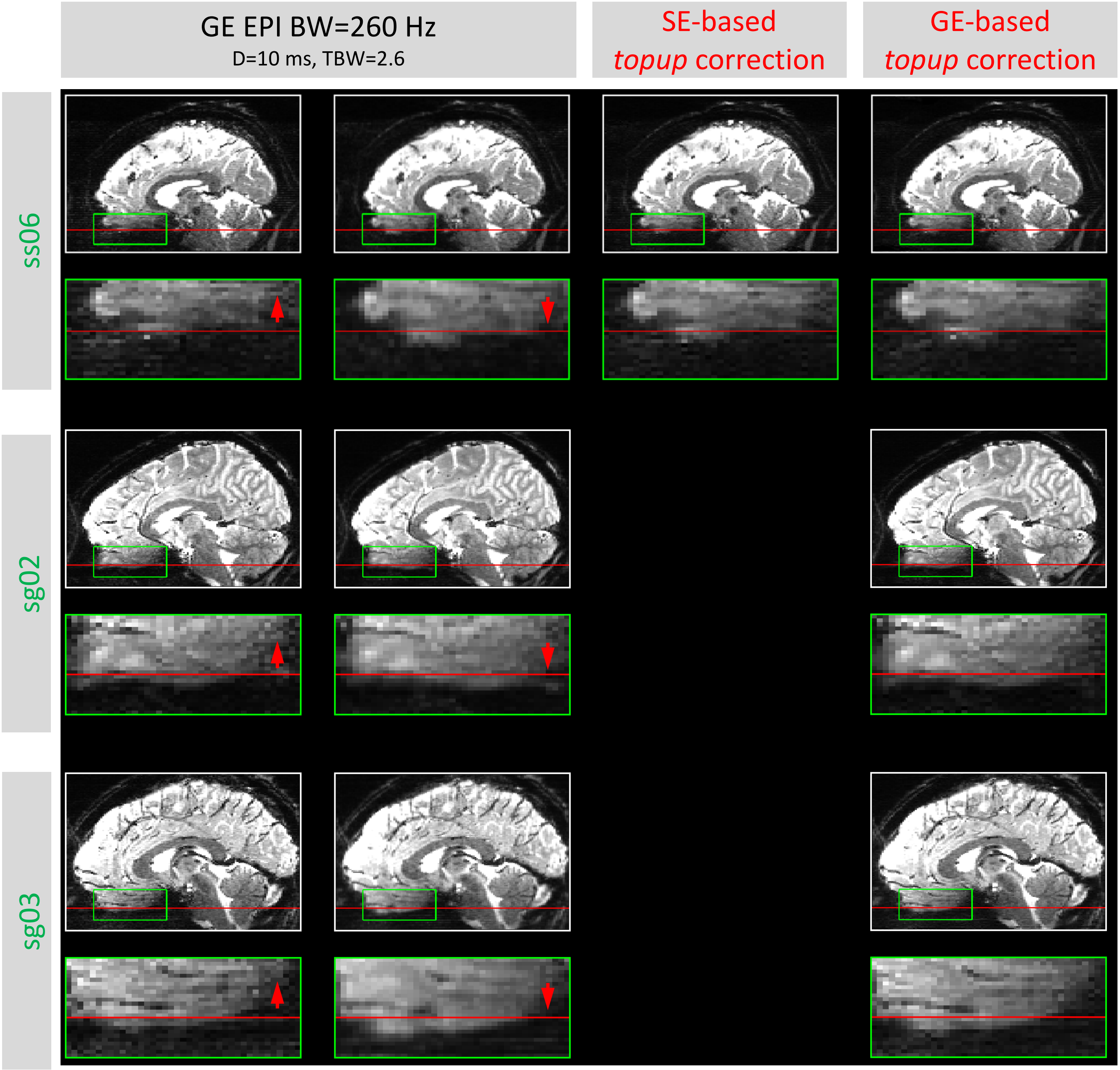


**Supplementary Figure 1 continued.**

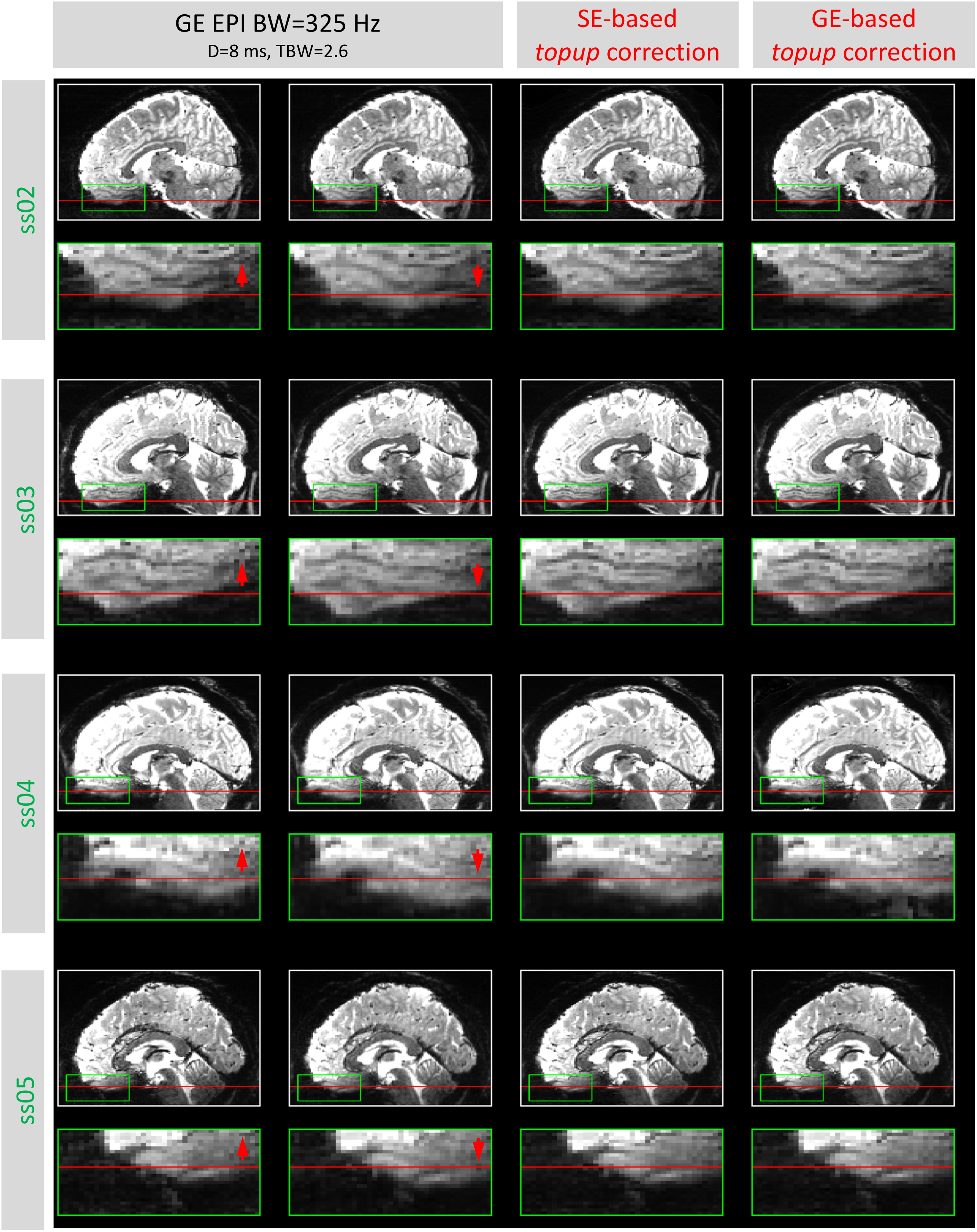
GE EPI BW=325 Hz
D=8 ms, TBW=2.6
SE-based
topup correction
GE-based
topup correction
ss02
ss03
ss04
ss05

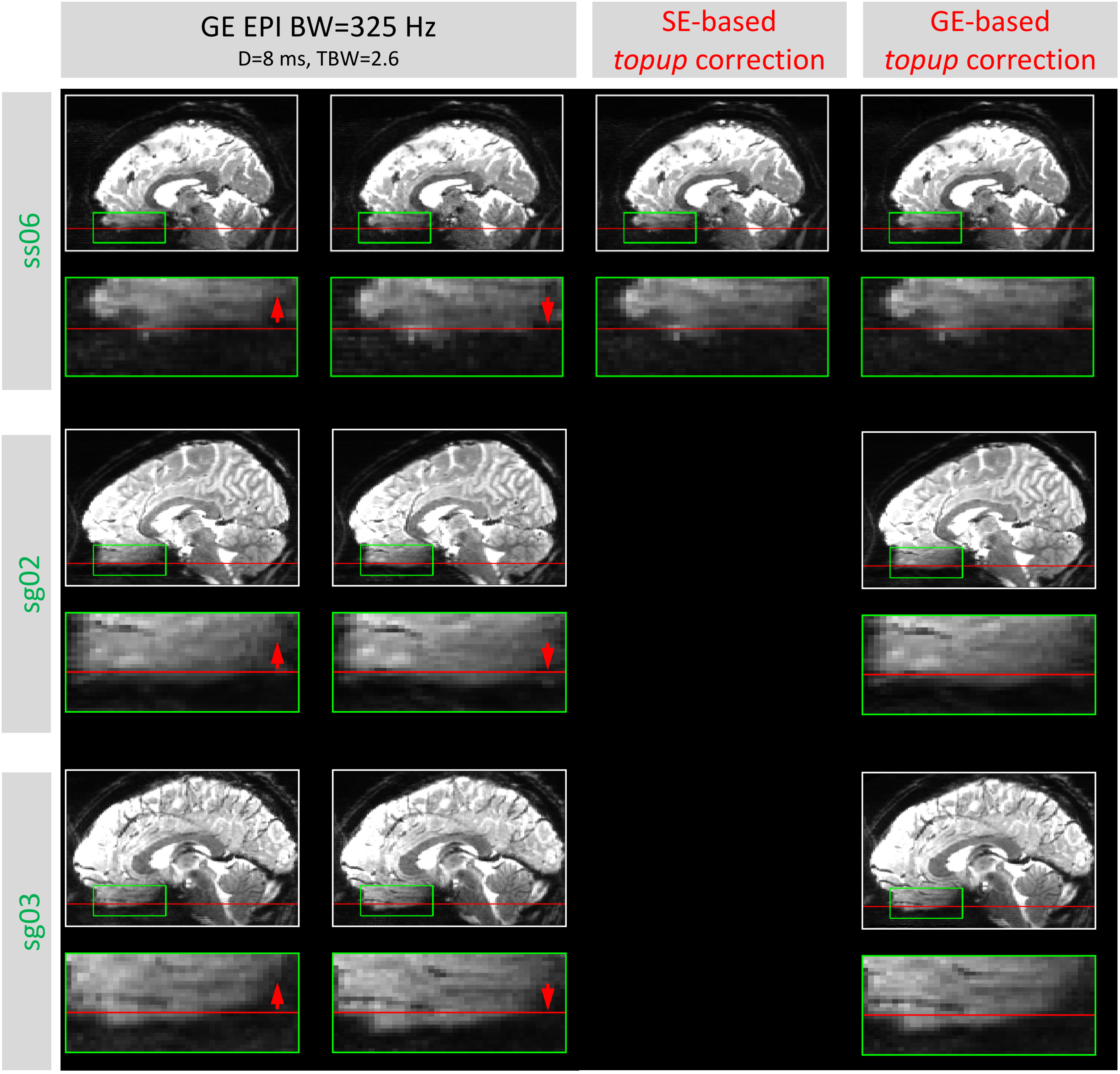


**Supplementary Figure 1 continued.**

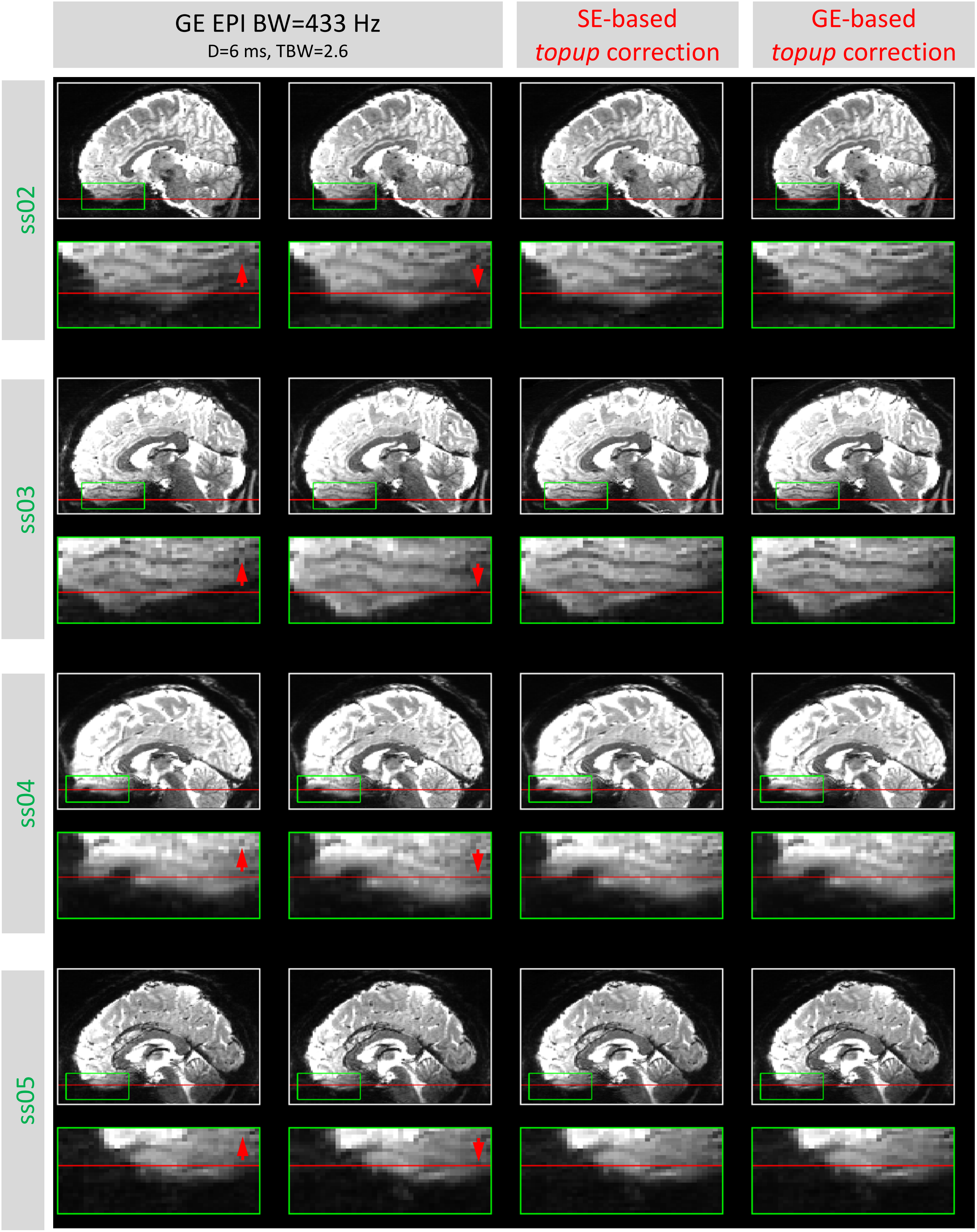
GE EPI BW=433 Hz
D=6 ms, TBW=2.6
SE-based
topup correction
GE-based
topup correction
ss02
ss03
ss04
ss05

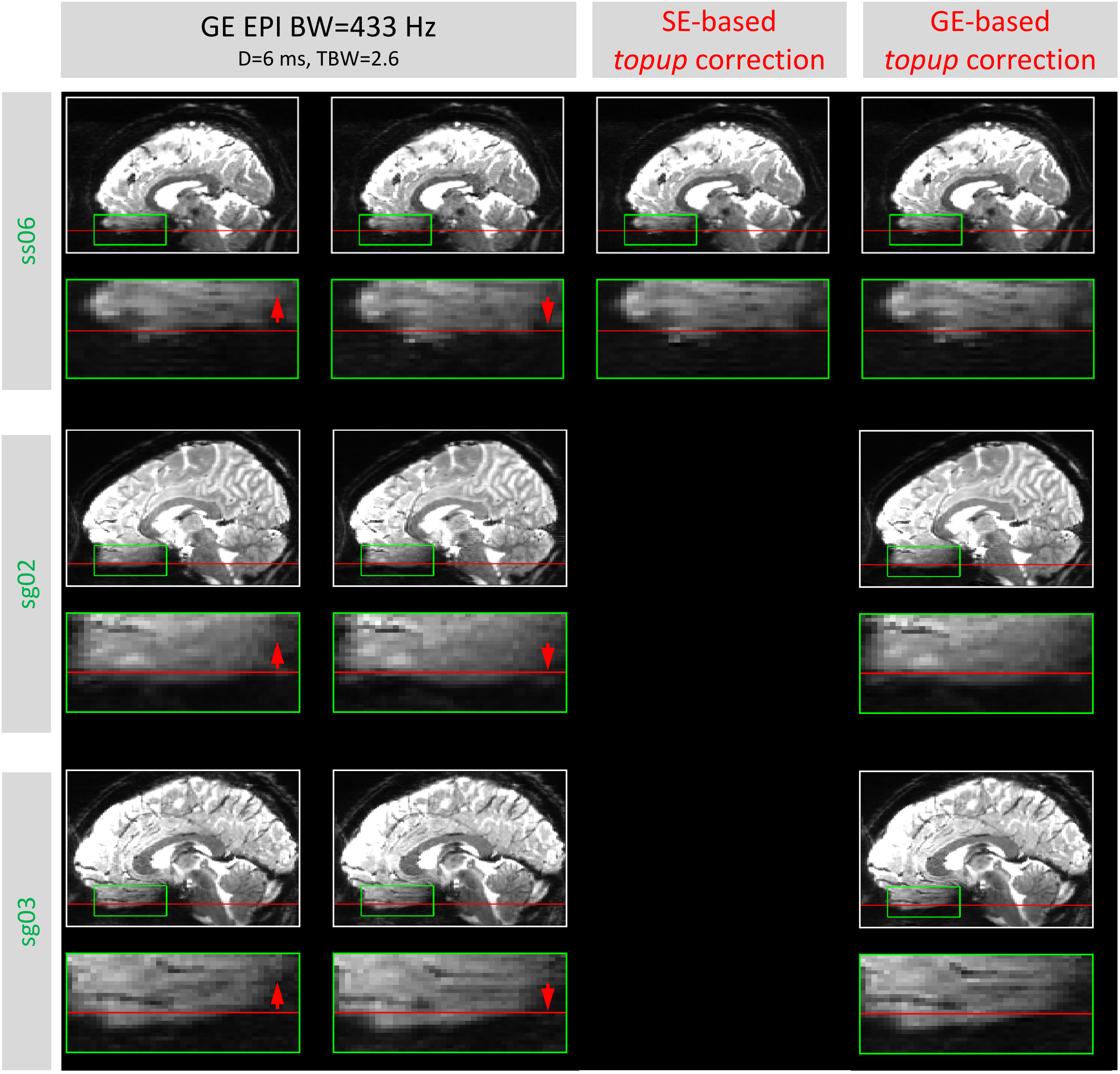


**Supplementary Figure 1 continued.**

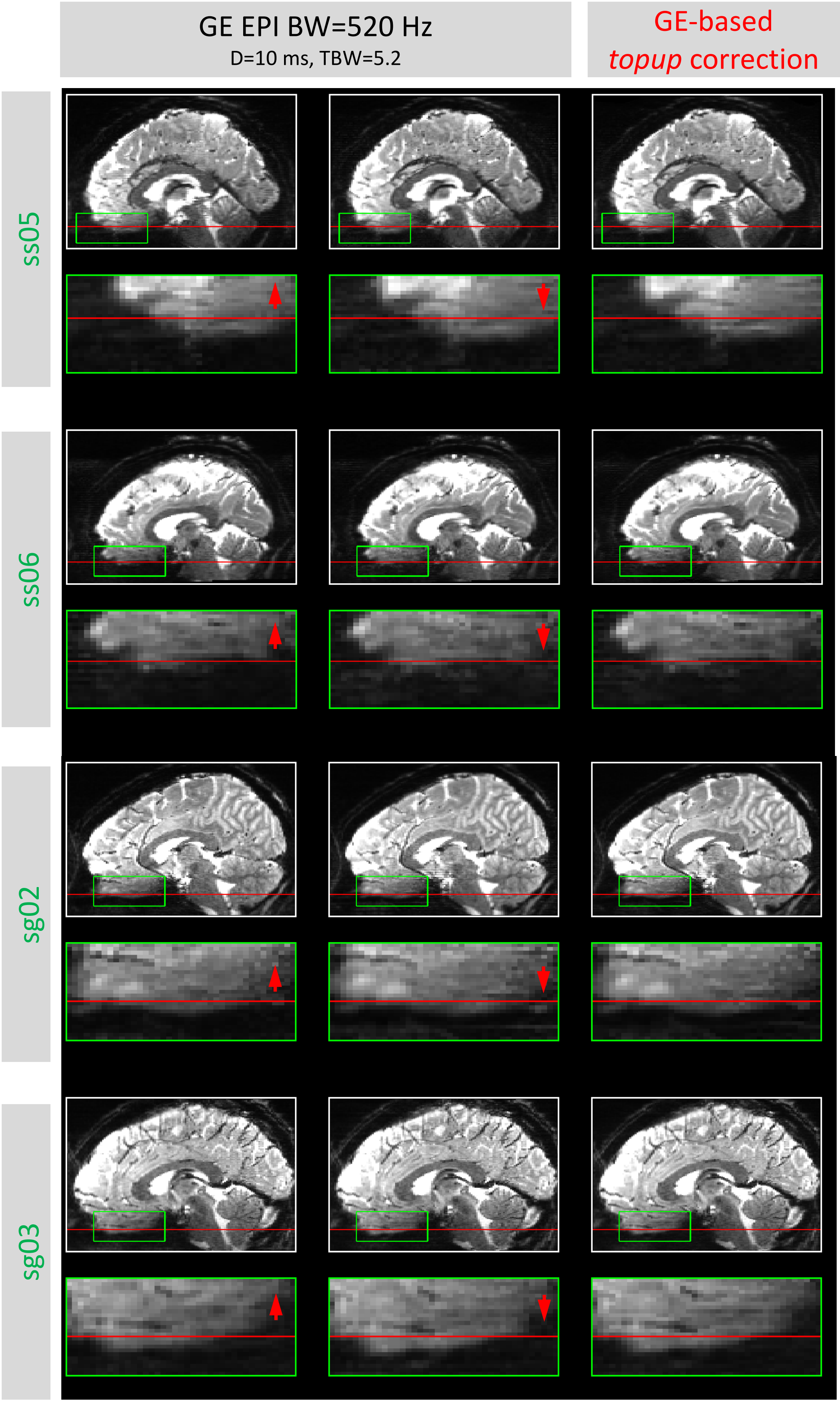


**Supplementary Figure 1 continued.**

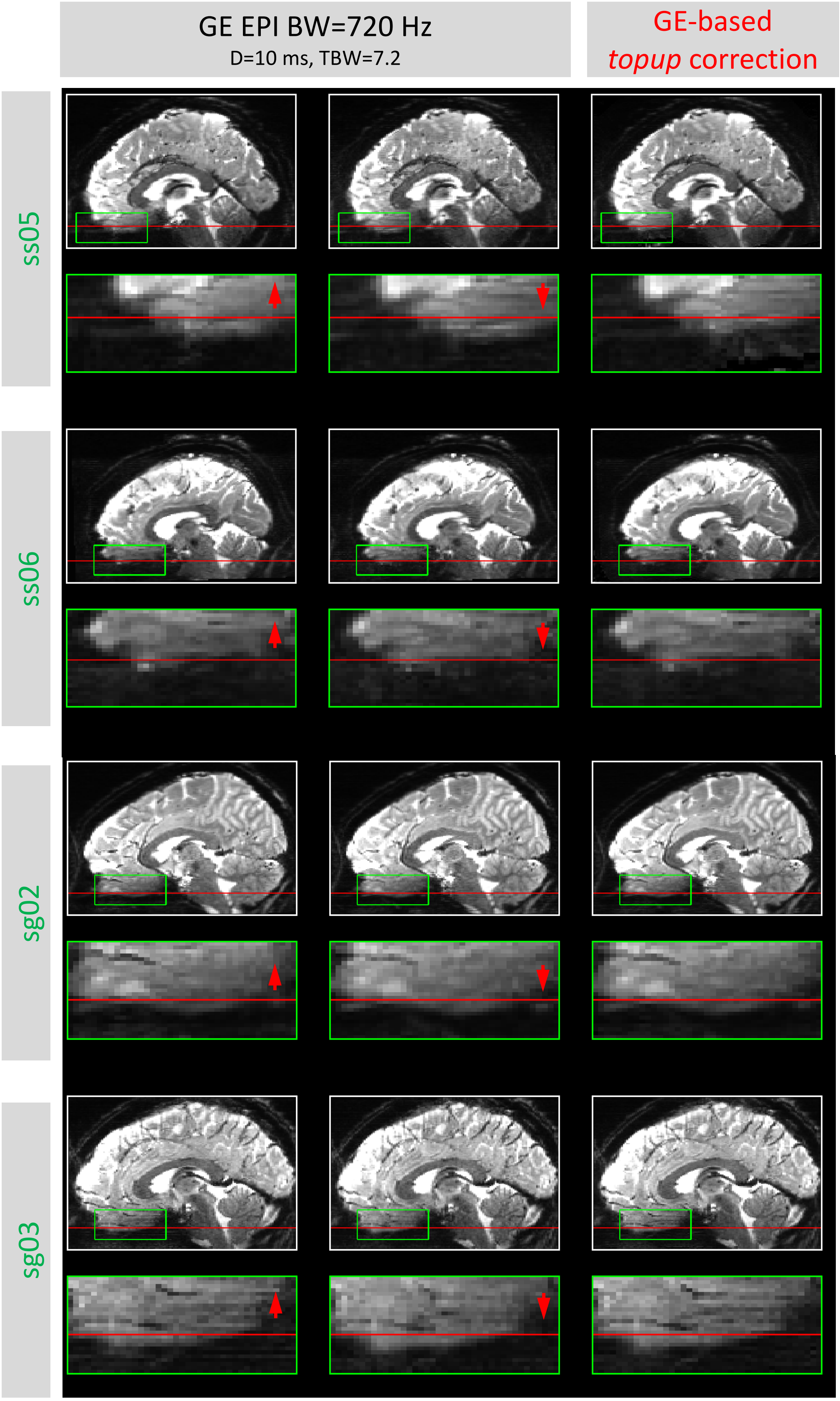


**Supplementary Figure 1 continued.**

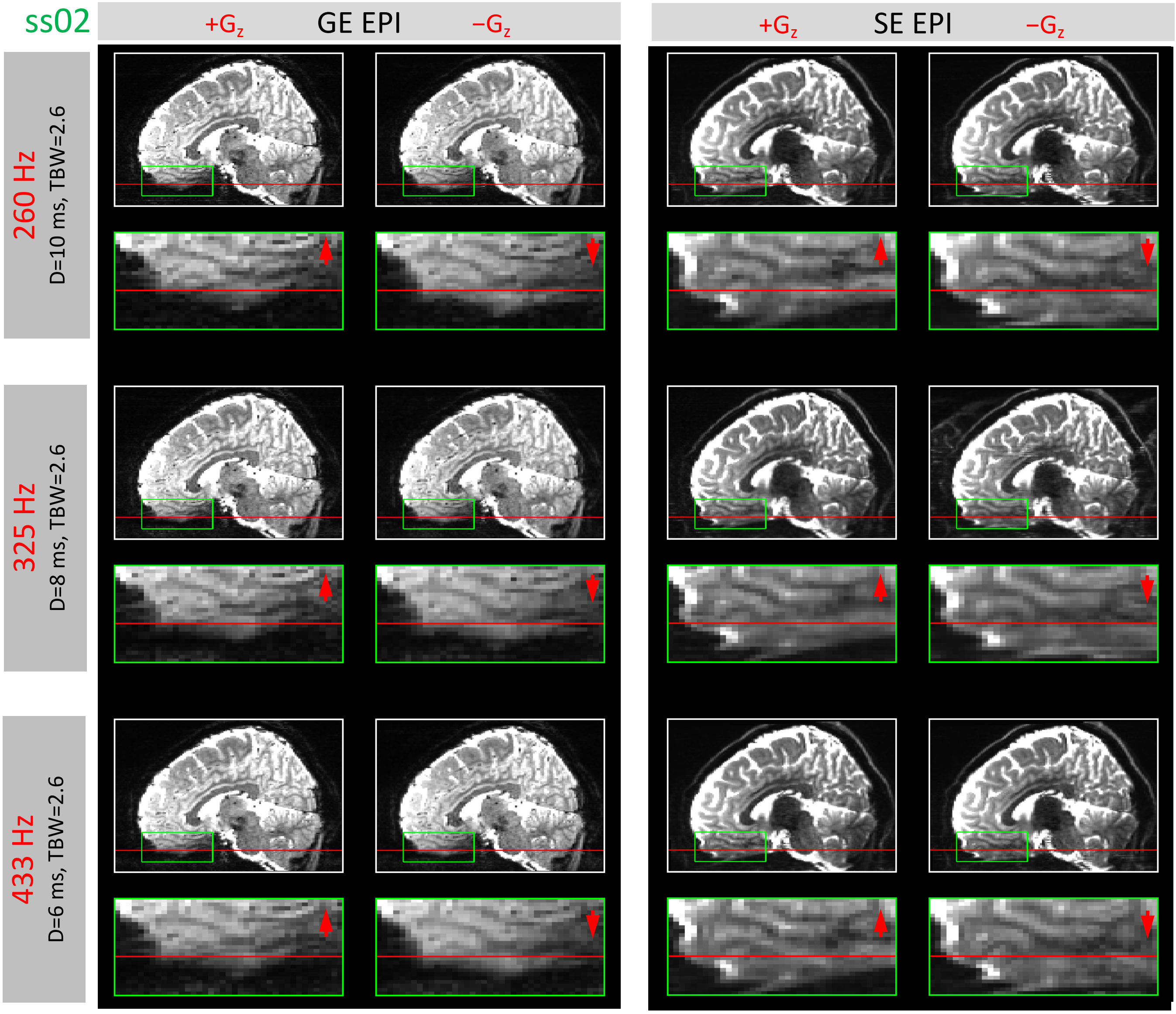


**Supplementary Figure 2.** Example sagittal reformats of gradient-echo (GE, left) and spin-echo (SE, right) EPI data from a representative subject, with 1-mm slice thickness and 1.5-mm in-plane resolution; positive (+$G_z$, left) and negative (−$G_z$, right) slice-encoding direction is indicated by red arrows. Rows are ordered according to increasing slice-encoding BW values achieved by changing either the slice-select/slice-refocus pulse duration (*D*) or time-bandwidth product (TBW); see text for details. As in Figure 3, the horizontal red lines are included in the same position in each pair of images to help visualize shift of voxels due to distortion; magnified inserts correspond to a region indicated by the green box where distortion is pronounced.

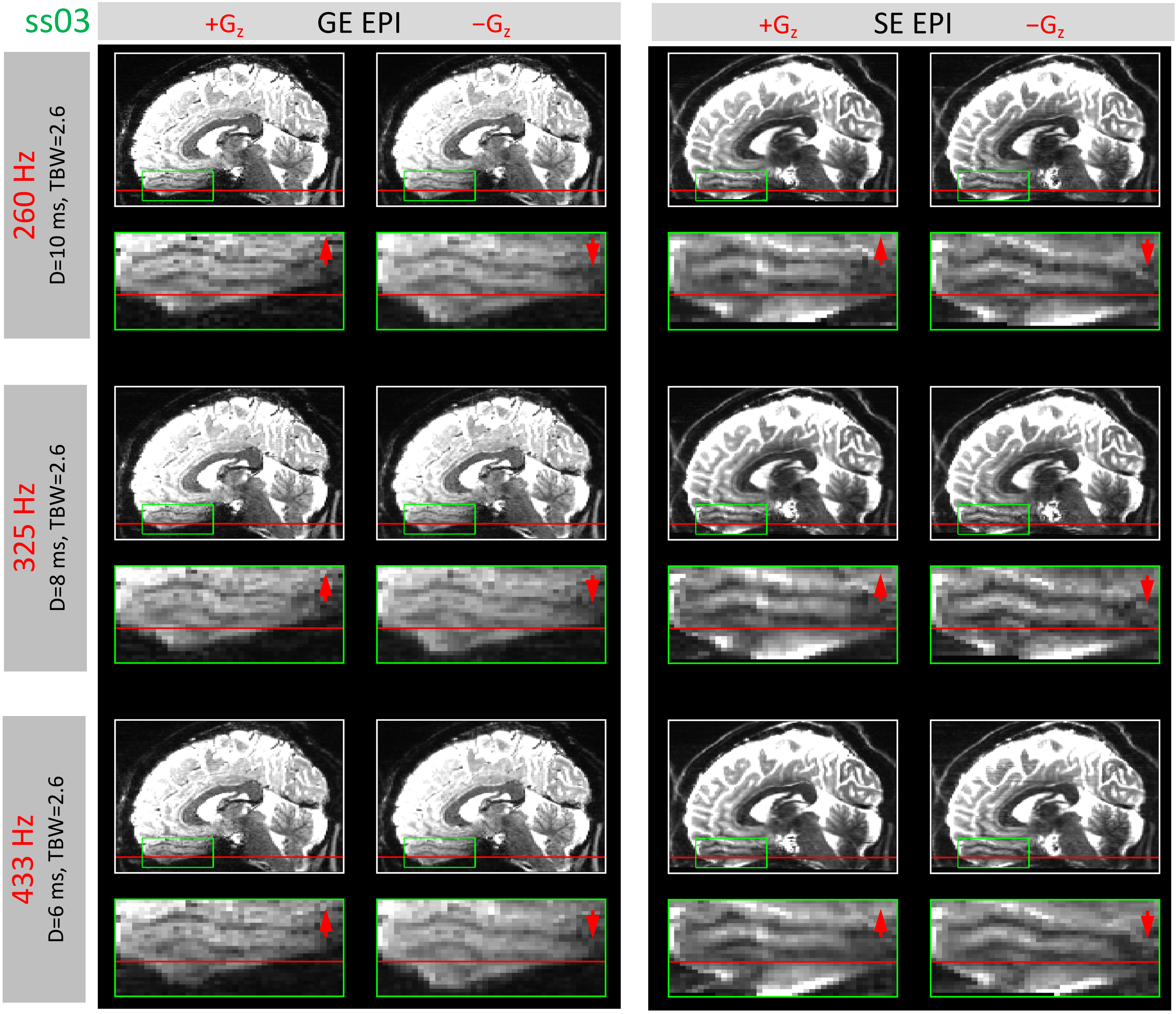


**Supplementary Figure 2 continued.**

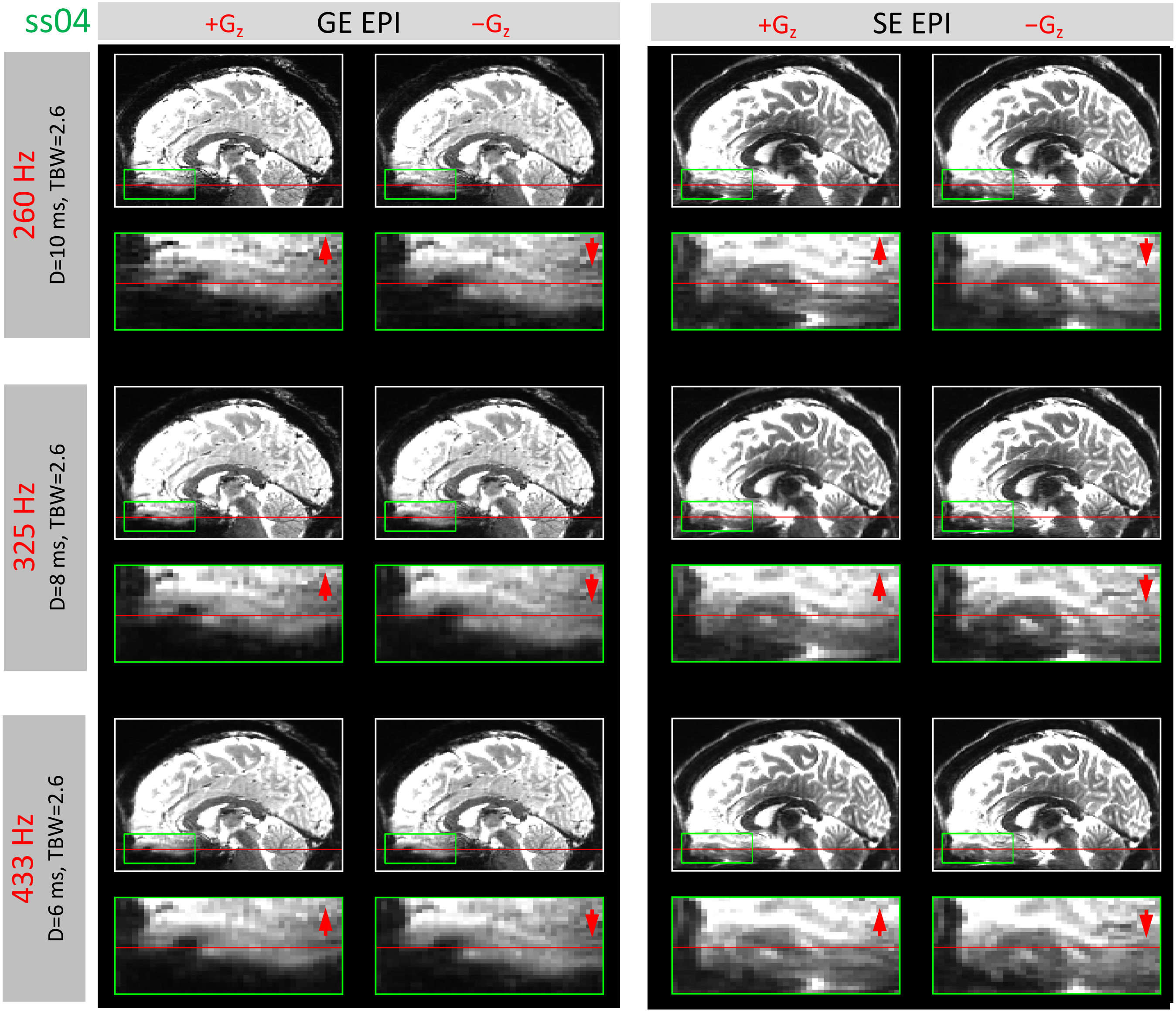


**Supplementary Figure 2 continued.**

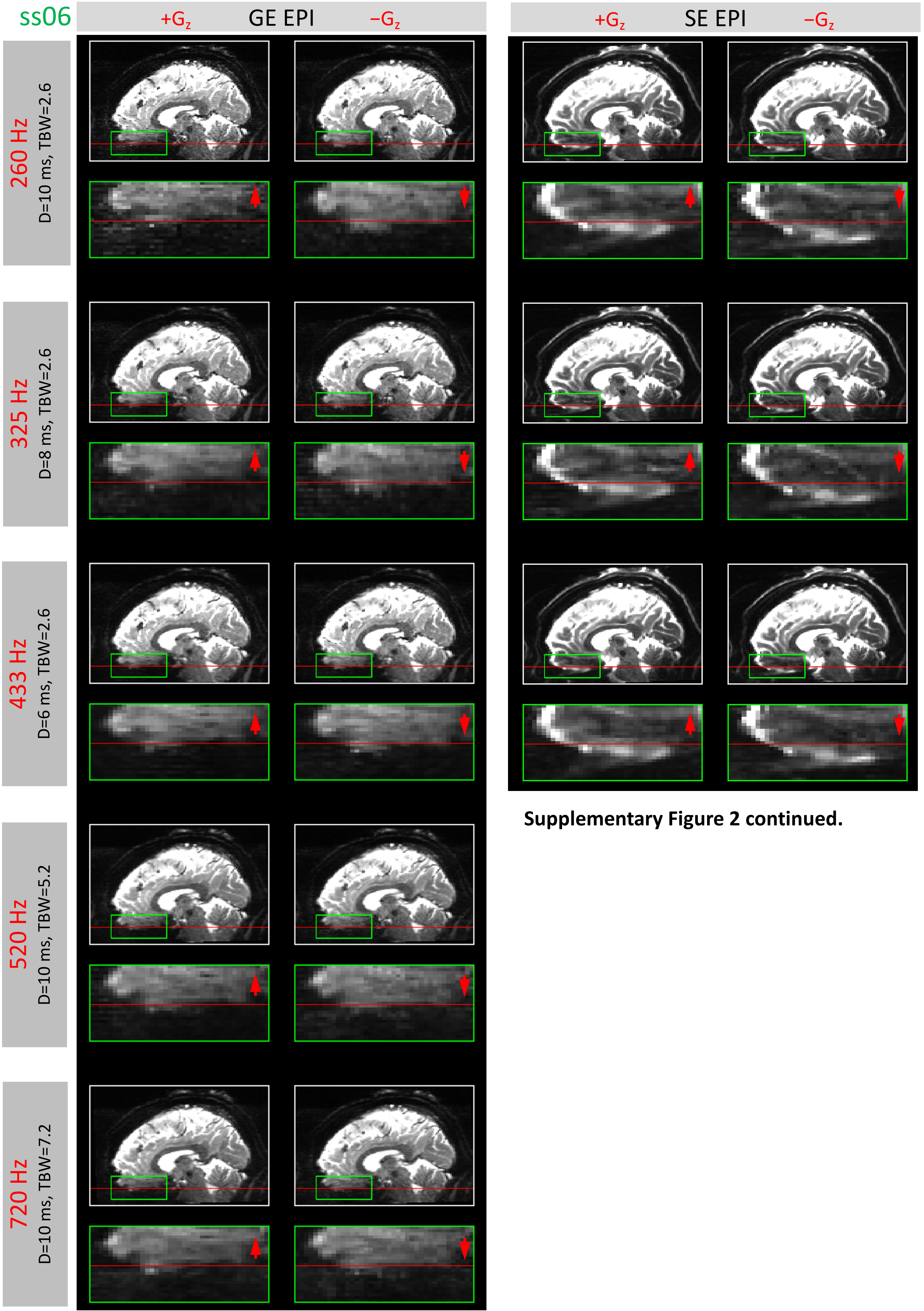


**Supplementary Figure 2 continued.**

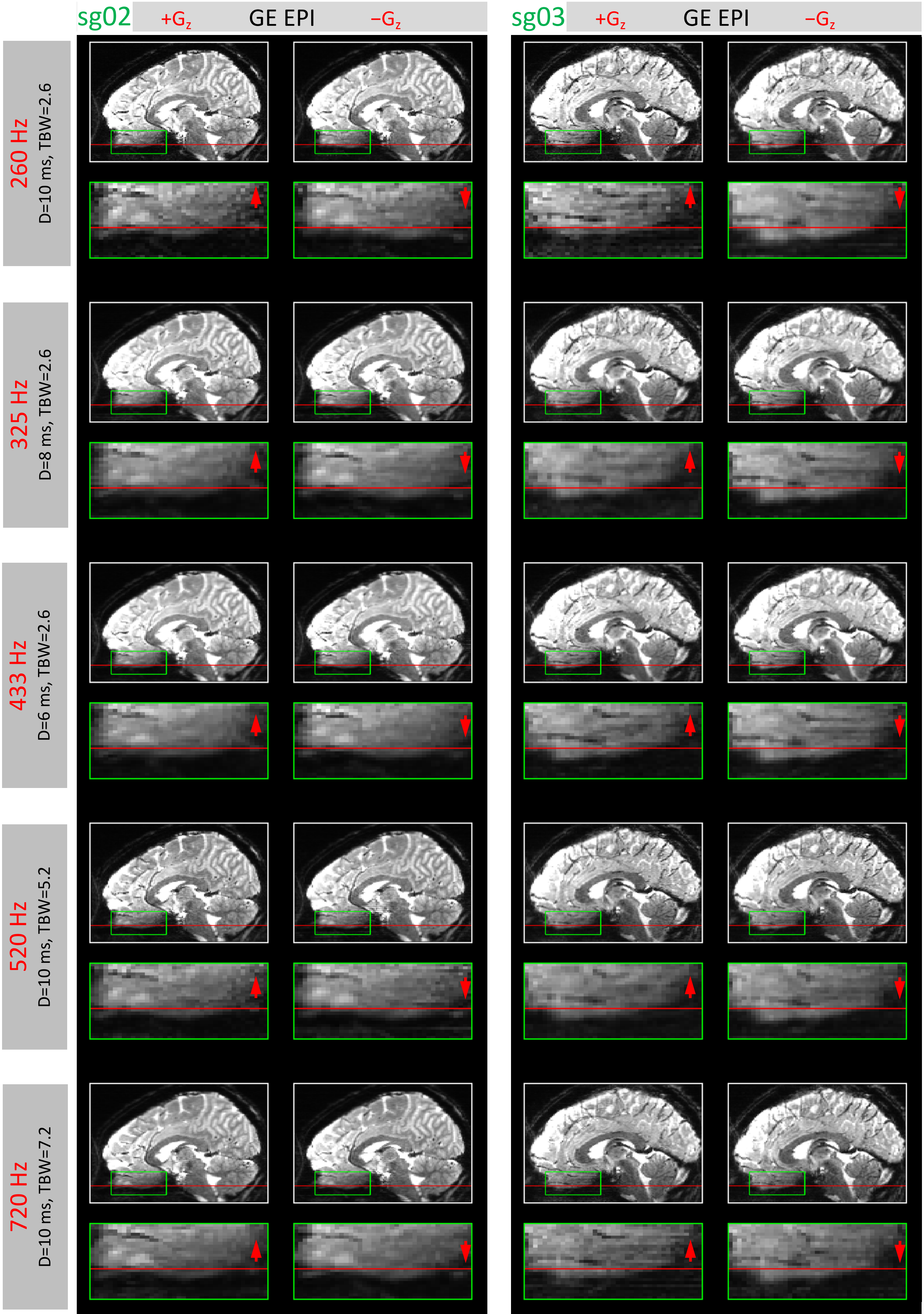


**Supplementary figure 2 continued.**
